# Impaired Regulation of Redox Transcriptome during the Differentiation of *i*PSCs into *Induced* Cardiomyocytes (*i*CMs)

**DOI:** 10.1101/519793

**Authors:** Gobinath Shanmugam, David Crossman, Johnson Rajasingh, Brian Dalley, Jianyi Zhang, Namakkal S. Rajasekaran

## Abstract

**Background:** Reprogramming of somatic cells into pluripotent stem cells (*i*PSC) and subsequent differentiation into *i*PSC-derived cardiomyocytes (*i*CM) seems to be a promising strategy for cardiac regenerative therapy. However, recent failure or poor outcomes in cardiac cell therapy warrants further investigation focusing on the infarction/wound environment (site of healing) to improve the cardiac regenerative medicine. Here, using next generation sequencing (NGS), we analyzed the global transcriptome to discover the unidentified genes/pathways that are crucial for cell survival, cytoprotection and mitochondrial dynamics during the differentiation of iPSC into iCM.

**Methods:** High throughput NGS was performed RNA from human *i*PSCs and *i*CMs (n=3/group) and analyzed the global changes in the transcriptome during differentiation. Furthermore, Ingenuity Pathway Analysis (IPA) and Gene Ontology (GO) for biological process were performed to understand the transcriptional networks that are involved during *i*CM differentiation. RNA-seq data were further validated by qRT-PCR analyses.

**Results:** Global transcriptome analysis revealed that ~9,290 genes (log_2_ FC >2) were significantly altered in human *i*CMs compared to the parent *i*PSCs, in which 4,784 transcripts were substantially upregulated and 4,506 transcripts were down-regulated during differentiation. GO enrichment and IPA analyses revealed the top 10 regulatory networks (i.e. hierarchical order) involved in differentiation of *i*CMs including cardiomyocyte remodeling, integrin-linked kinase signaling, Rho family of GTPases, etc. Surprisingly, none of the top 10 pathways listed the genes liable for redox signaling networks that are crucial for the basal cellular redox homeostasis, Nrf2-dependent antioxidant defense, mitochondrial functions and cell survival. Our deeper and unbiased analysis of this data revealed that the genes involved in above canonical signaling pathways are found in the middle of the inverted vertical cone. Of note, although these pathways are significantly altered during the differentiation (of iPS into cardiomyocytes), a majority of them are ranked low in the hierarchical list (>150). Validation of the randomly selected genes representing various pathways real-time qPCR confirmed the global transcriptome changes observed in NGS.

**Conclusion:** We highlight the significance of Nrf2-redox and mitochondrial transcriptome during differentiation of iPSC into iCMs. Thus, targeting the redox signaling mechanisms in iCMs may enhance their efficiency for cell therapy and improved myocardial repair.

## Introduction

Human induced pluripotent stem cells (hiPSCs) have the ability to proliferate and differentiate into other functional cell types including beating cardiomyocytes (induced cardiomyocytes; *i*CM) (1-3). The cardiac lineage represents a possible source for myocardial regenerative therapy (4, 5). These iCMs also provide a reliable platform for researchers to study the disease mechanisms, drug screening and gene specific gain and loss of functions (6-9) at the cellular level. A successful differentiation of *i*CMs and their faithful recapitulation of cardiomyocyte structure and function including contraction, electrophysiological signaling properties, and dynamic metabolic capacity to combat various stress in the infarcted site and repair the infarcted human myocardium (8). Thus far, studies mainly focused on signaling pathways that are crucial for cardiac differentiation and maturation (2) rather the mechanisms for cellular stability and sustainability against various stress conditions including oxidative & inflammatory stresses (13, 14). During the repair, following infarction, the human myocardium exhibits a severe metabolic stress (i.e. switches to lipid oxidation) due to increased inflammation and reactive chemical species (RCS) production (i.e. oxidative milieu) (15). The mounting stress on aging further induces RCS leading to a hyper-oxidative condition in the infarcted regions of the myocardium (16, 17). Currently, transplantation of iPS-derived cardiomyocytes (iPS-CMs) into the infarcted areas results in their death due to necrosis and/or apoptosis, thereby reducing their survival and poor implantation/repair (18-20). Therefore, understanding the redox-dependent mechanisms that are vital for augmented survival and engraftment of iPS-CMs is essential.

For a better understanding of the redox status of iCMs, we performed genomic and bioinformatics analyses in human iPS (hiPS) and hiPS-derived cardiomyocytes (hiPS-iCM) to identify novel genes and pathways that are associated with the redox transcriptome. Most of the published reports focused mainly on analyzing the pathways involved in reprogramming, differentiation, maturation and metabolic signaling of induced cardiomyocytes (1, 2, 8, 21, 22). Outcomes of these transcriptome analyses included genes that are dramatically changed (differential expression >50 fold) in iCM versus iPS cells, but ignored several thousands of genes that are significantly changed at >2.0 to 20 fold resulting in masking the significance of several other key pathways during differentiation. Therefore, a closer analysis of genes that are significantly altered (even at 2 or more-fold) will provide a better understanding of other key pathways in the context of redox signaling including Nrf2-redox, antioxidant, hypoxia signaling and mitochondrial energy dynamics during differentiation of iCM. Moreover, most of the previous reports discussed the gain of function (upregulated genes) mechanisms but left the downregulated genes that are associated with loss-of-function events. This gap in knowledge is expected to be associated with a huge failure in cell-based therapy for myocardial repair.

Under normal/unstressed conditions, RCS play a key role in cellular physiological signaling and regulate normal cellular processes such as induction of proliferation, cell-cycle, and differentiation (23, 24). However, an excessive accumulation of RCS induces oxidative stress leading to apoptotic, necrotic and fibrotic events (25, 26). In fact, RCS signaling is necessary for the differentiation of iPS into iCM (23, 27), but a chronic increase in RCS is detrimental to these cells as it could impair both the proliferation and differentiation processes (24, 25). Real time quenching of RCS is vital to maintain redox homeostasis, which is often controlled through multiple biochemical events. These pathways include Nrf2-mediated oxidative stress response, nitric oxide & hypoxia signaling, mitochondrial bioenergetic dynamics, etc. (28-31). Of note, the redox levels and pathways involved in redox signaling have not been thoroughly studied in the differentiating iPS-iCMs.

To address this important gap in knowledge, we performed next generation RNA sequencing (NGS-RNAseq) using RNA from h*i*PSCs (proliferating/undifferentiated) and h*i*CMs (differentiated) and determined their transcriptome profiles. Deep sequencing results indicate that the *i*CMs showed 9,290 genes significantly (>2.0 fold) altered transcripts that are previously unidentified. Furthermore, Ingenuity Pathway Analysis (IPA) was performed for the top 2,000 significantly changed genes and individual IPA analysis for genes that are changed above 100, 50, 10, 5 and 2-fold in iCMs vs. iPS cells to elucidate previously uncovered pathways and their regulatory networks responding to the differentiation process.

## Methods

### Reagents

All cell culture reagents were procured from StemCell Technologies. Primers for qPCR were designed using Primer Bank website (https://pga.mgh.harvard.edu/primerbank/) and verified by Primer-Blast. All primers were purchased from Eurofins Genomics, Louisville, KY. RNeasy kit, reverse transcription kit, and QuantiTect SYBR Green PCR kit were purchased from Qiagen Inc., Valencia, CA. All other chemicals including *RNAlater* and ethanols were purchased from Sigma-Aldrich unless otherwise stated.

### Induced Pluripotent Stem cells and differentiation into cardiomyocytes

All cells were maintained in a sterile condition at a constant temperature (37 °C) in a Heracell 150i CO_2_ Incubator with 5.0% CO_2_. Both induced Pluripotent Stem cells (iPSCs) and induced Cardiomyocytes were cultured in Matrigel coated plates. iPSCs were cultured in mTesR1 Basal medium supplemented with mTesR1 supplements. The iPSC culture medium was replaced every 48 hours. Once cells reached the 95% confluency, iPS cells were initiated to differentiate into cardiomyocyte by adding cardiomyocyte differentiation supplement A and after 48 h (day-2), supplement B was added. After 48hours of incubation with supplement B (day-4), cells were induced with Supplement C for 48 hours (day-6) and again supplement C was added for another 48 hours (day 8). At the end of the last 96 hours of incubation with supplement C, cells started to show cardiomyocyte properties like beating cardiomyocytes.

Further induced cardiomyocytes were cultured for another 2 days with cardiac maintenance medium. At day 10, induced cardiomyocytes were harvested for RNA isolation (21, 32).

### Isolation of RNA and Next generation mRNA sequencing

RNA was isolated from Induced Pluripotent Stem cells (iPSC), and iPS-derived cardiomyocytes using RNeasy mini kit (Qiagen, 74106). Cells were lysed using RLT lysis buffer and total RNA was isolated (33). RNA concentration and purity was measured using Thermofisher Nanodrop One^c^. After confirmation of RNA purity, using oligo(dT) magnetic beads intact poly(A) transcripts were purified from total RNA and mRNA sequencing libraries were prepared with the TruSeq Stranded mRNA Library Preparation Kit (Illumina, RS-122-2101, RS- 122-2102). The detailed protocol for RNA sequencing was discussed in earlier (34, 35).

### Ingenuity pathway analysis

All differentially expressed genes obtained between iPSCs vs iCM cells from RNA sequence analysis were subjected to Ingenuity pathway analysis. Based on the fold changes in mRNA expression, five different IPA (>100, >50, >10, >5 and >2) runs were performed relative to the iPSCs expression profile. Log-transformed p-values of significantly enriched canonical pathways associated with the iPSC-derived cardiomyocytes (iCMs) were used to construct a heat maps. IPA analysis included gene lists for each biological process, and the top 15 up and down regulated DEGs in iCM cells within each major biological function category were assembled into heat maps based on their log2 relative expression ratios (34).

### Gene Ontology and String analysis

Gene Ontology for biological process (version 2018) was performed for upregulated, down-regulated and top most differentially regulated genes individually using *Enrichr* online tools to determine the significant top changes in the biological process during cardiomyocyte differentiation. Further, to investigate the role of the differentially expressed genes during cardiomyocyte differentiation, we performed a protein-protein interaction analysis to identify and recognize functional elements and key proteins among the differentially regulated genes in each pathway. Online STRING tool (https://string-db.org/) with medium confidence score (0.400) was used for the network analysis and excluded interactions based on text mining (35, 36).

### Real-time qPCR analysis

RNA sequencing results were validated using real time qPCR analysis. cDNA was synthesized using 1.25 μg of RNA with QuantiTect reverse transcription kit (Qiagen, 205313). 25-50ng cDNA and 10 pmol gene specific primer were used in a 10 μl SYBR green reaction mix (Qiagen, 204056) for Quantitative RT-PCR (Supplemental Table 1) and amplified in a Roche Light Cycler 480 (Roche, Basel, Switzerland). Relative expression was quantified using Ct values, and expression fold-change was calculated by normalization to the Ct of housekeeping genes *GAPDH* according to the 2^−ΔΔCt^ methods (37, 38).

### Statistical analysis

All data are represented as mean ± SEM. Student’s t-test was used for comparisons in iPSCs vs iCMs. All analyses were performed using GraphPad Prism 7. Differences were considered significant at the values of *p< 0.05, **p< 0.01, and ***p< 0.001.

## Results

### Generation of h*i*CM from h*i*PSC cells of the same origin

To reduce line-to-line variations and genetic backgrounds, h*i*PSCs were purchased from Stem Cell Technologies. Before performing the RNAseq analysis, two 6-well plates of cells from the same line were cultured. One plate was collected for iPSC-RNA and the second plate was differentiated into cardiomyocytes. After differentiation, iCMs cells were further cultured to observe the contractile phenotypes. RNA from differentiated beating cardiomyocytes (iCMs) was used for RNA seq analysis for the *i*CM group. Before collection of RNA, morphology of *i*PSCs and *i*CMs was recorded by light microscopy (Figure 1A). To examine the consistency within the data, we used hierarchical clustering of all samples used for next generation RNA sequencing (NGS) data, which displayed a distinctive grouping in accordance with their respective groups (Figure 1B). These high through put sequencing analysis using RNA-seq from h*i*PSC and h*i*CMs detected ~26,194 genes and approximately ~9,290 genes (~35.08%, >1.5) were significantly altered in h*i*CMs cells. The hierarchical clustering for ~9,290 genes was represented as a heatmap (Figure 1C). From these differentially expressed genes, 52.1% (~4,784) transcripts were upregulated and 47.9% (~4,406) transcripts were down-regulated during induced cardiomyocyte differentiation. Overall, the Venn diagram described that 2.4% transcripts were highly expressed (> 100 fold), 4.7 % transcripts were greatly altered (> 50 fold), 20 % transcripts were prominently altered (>10 fold) and 35.5% transcripts were moderately changed (>5 fold) (Figure 1D).

**Figure 1.**
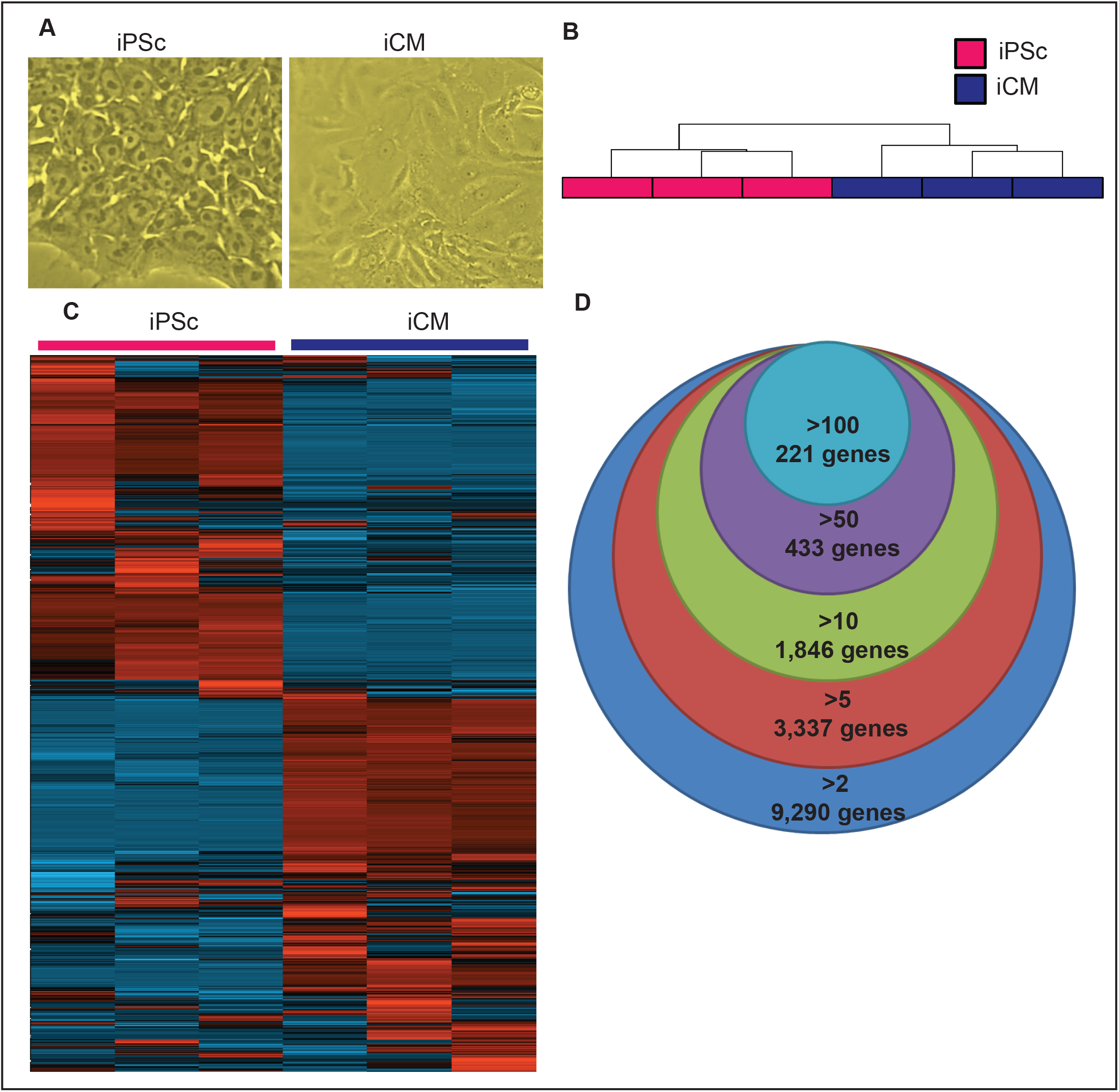
Generation and Comparison of *i*PSCs and *i*CMs. A). Light microscopy images revealing the morphology of induced pluripotent stem cells (*/*PSs) and their differentiated induced cardiomyocytes (*i*CMs). B). Hierarchical clustering of normalized FPKM values from RNA-seq revealing distinctive grouping of 3 samples from each group. C). Heat map demonstrating 9, 290 differentially expressed genes (of more than 2FC) in *i*CMs compared to *i*PSs. D). Venn diagram representing the number of genes significantly changed in induced cardiomyocytes from NGS RNA-seq compared to induced pluripotent stem cells with fold changes ranging from 2 to more than 100.

### Top altered canonical pathways and biological functions in iPS-derived cardiomyocytes using gene ontology

IPA was performed on the differentially expressed genes (DEG) based on their changes for above 100, 50, 10, 5- and 2-fold to explicate pathways and their regulatory networks involved in differentiation of *i*CM cells. The top pathways enriched in *i*CMs versus *i*PSC cells on biological processes were expectedly related to Cardiomyocytes remodeling, Integrin-linked kinase signaling, Rho family of GTPases which is involved in organelle development, cytoskeletal dynamics (Table-1). To further investigate the differences between *i*PSC and *i*CMs, we performed gene ontology (GOs) for the altered transcripts and GO results displayed the biological process highly related to cardiomyocyte/cardio myoblast differentiation. Next we performed Gene Ontology for biological process using upregulated and down-regulated genes separately, the topmost outcome of biological process for upregulated genes being related to DNA replication and chromatin remodeling (Figure 2A). The biological process for down-regulated genes is largely related to hormonal responses and unfolded protein responses (Figure 2B). Further GOs for highest upregulated genes showed cardiac development and differentiation functions including two functions related to insulin signaling, three functions related to cardiac development, and one each for renal sodium excretion, spleen development etc., (Figure 2C).

**Figure 2.**
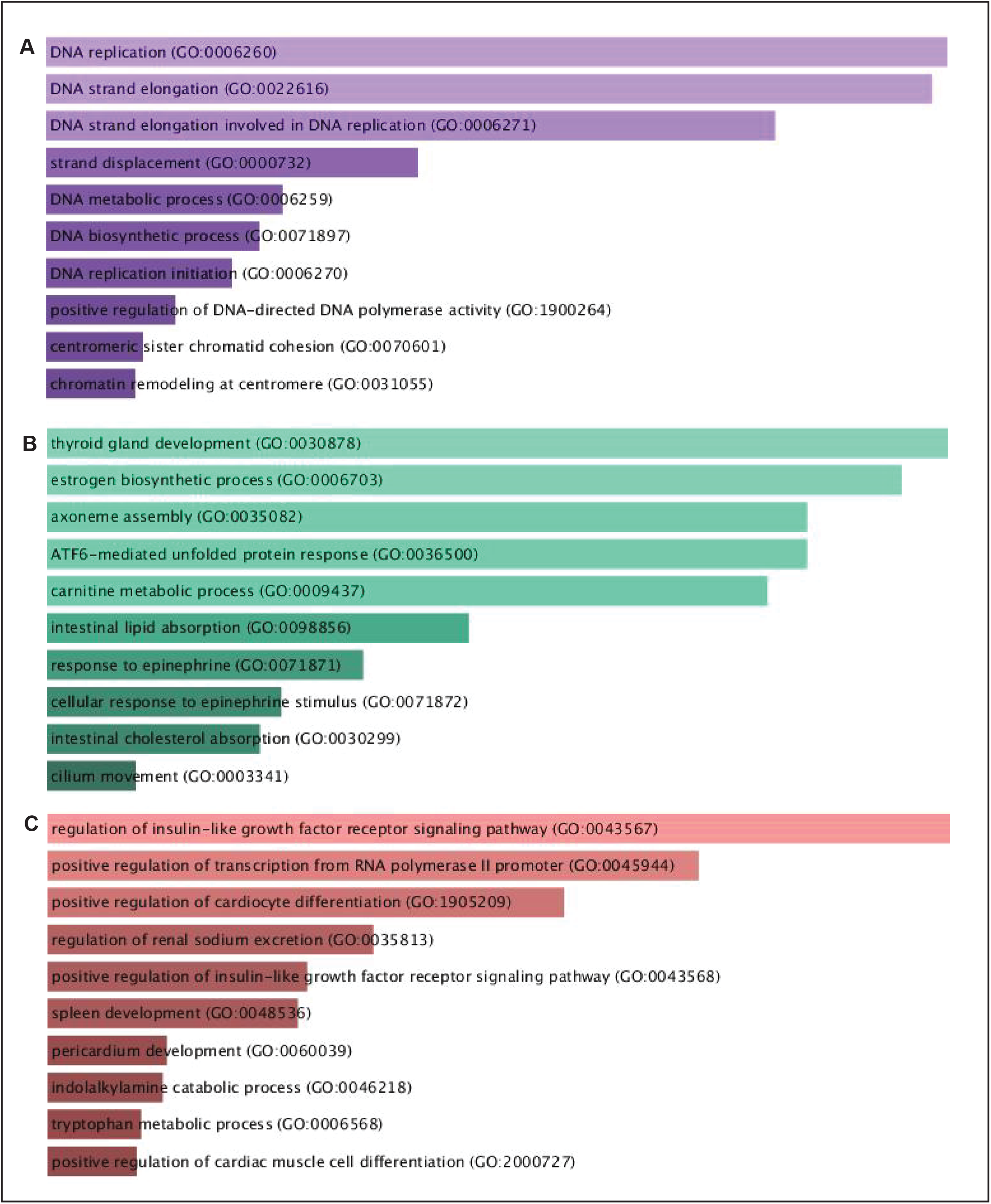
Significantly altered Gene Ontology (GO) terms of biological process in *i*PSC-derived *i*CMs. A). Gene Ontology (GO) terms of biological process enriched functional categories for significantly upregulated genes above 2 fold, p<0.05. **B)**. Gene Ontology (GO) terms of biological process enriched functional categories for significantly down-regulated genes above 2 fold, p<0.05. **C)**. Gene Ontology (GO) terms of biological process for top differentially expressed genes above 100-fold using RNA-seq.

### Pathways involved in redox signaling

To further elucidate the expression patterns of redox signaling pathway genes during *i*CM differentiation, we selected some unique pathways that are potentially involved in redox regulation such as Nrf2-glutathione mediated oxidative stress response, hypoxia signaling, Nitric-oxide signaling in the cardiovascular system, mitochondrial dynamics, Antioxidant action of Vitamin C and Protein Ubiquitination (Figure 3-5). Additionally, based on gene ontology we have performed mRNA-mRNA interaction studies and prepared networks based on their interactions (Figure S1-S6). These genes selected from canonical pathways were further studied within their expression patterns during differentiation. The top 30 (15 each of up and downregulated) DEGs within these major biological functions altered during cardiomyocyte differentiation were compared to their relative expression in *i*PSC cells, and 6 of each up and down-regulated genes were chosen for real-time qPCR validation from each pathway. Most significant correlations were observed for most of the genes between RNA-seq derived fold expression ratios and log_2_ transformed mean fold-change observed in validation experiment.

**Figure 3.**
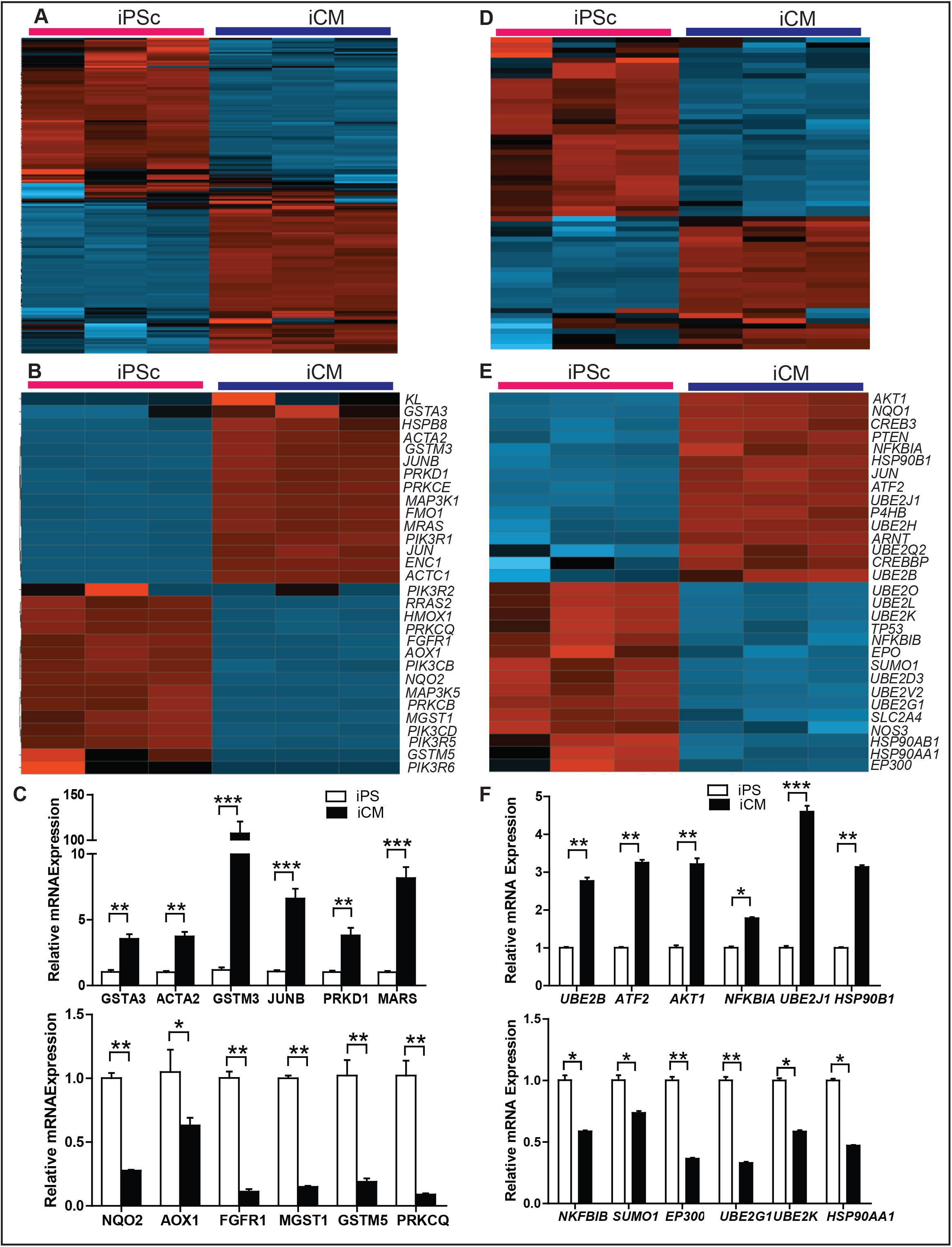
Significantly altered Canonical pathways in biological relevance with redox signaling. A). Hierarchical clustering (heat map) showed differential expression of all genes involved in Nrf2-Glutathione mediated oxidative stress response pathway identified by Ingenuity Pathway Analysis. **B)**. Top 15 up and down regulated DEGs in Nrf2-Glutathione mediated oxidative stress response pathway represented by heat map. **D)**. Differentially expressed genes for all genes involved in hypoxia signaling represented by heat maps using RNA-seq. **E)**. Top 15 up and down regulated DEGs in hypoxia signaling represented by heat map. **C and F)**. Real time qPCR validation of 6 up and down regulated genes from each respective pathway validated the significantly altered expression compared between *i*PSs vs *i*CMs cells. Gapdh was used as control (n=4-6/group). *p<0.05, **p<0.001, ***p<0.0001.

### Transcriptional analysis for Nrf2-mediated glutathione response and hypoxia transcriptomes during iPSC-cardiomyocyte differentiation

Nrf2 is a master regulator of antioxidant transcription and plays a major role in cytoprotective mechanisms (35, 37). During increased reactive oxygen species (ROS) production, Nrf2-medicated antioxidant pathway is triggered to compensate the effects of ROS (35, 37, 39). During iPS derived cardiomyocyte differentiation, an almost equal number of genes were differentially regulated (up and down regulated) in *i*CMs and this list of genes are represented as a heat map in Fig 3A-B. In Nrf2-mediated oxidative stress response pathway,180 genes were detected out of 230 genes known to constitute this pathway, whereas 58 genes were increased and 56 genes were decreased. From this list, the top 15 up and down-regulated genes were used to generate the heat-maps. A substantial list of genes were downregulated during cardiomyocyte differentiation including, *MGST1, GSTM5, PRKCQ, PRKCB* and *PIK3R*, while *PIK3R5ACTA2, GSTM3, FMO1, ACTC1* and *GSTA3* were increased during cardiomyocyte differentiation. RNA-seq data was validated using qPCR analysis showing *GSTA3, ACTA2, GSTM3, JUNB, PRKD1, MARS* gene expression was increased and *NQO2, AOX1, FGFR1, MGST1, GSTM5, PRKCQ* were down-regulated confirming the sequencing data. Protein-protein interactions were analyzed among the differentially expressed genes in Nrf2-mediated glutathione response pathway with standard parameter settings and higher interaction was observed in most of the proteins involved in this pathway (Figure S1).

Reduced oxygen condition activates the hypoxia signaling pathway (40) and hypoxia inducible factor (HIF) primarily regulates hypoxia mediated response (41, 42). IPA’s biological functions of hypoxia signaling in cardiovascular system involved many genes shown to protect the cells from hypoxia stress. In this particular pathway, 61 genes out of 75 mRNAs were detected and represented as a heat-map (Figure 3D). Further, the top 15 up and down-regulated genes were presented in a heat-map format for significant changes (Figure 3D-E). Among the 61 detected genes, 29 transcripts were significantly altered and the topmost up-regulated genes include *JUN, UBE2J1, NQO1, ATF2* and *P4HB* (>2.5), and down-regulated genes including *HSP90AA1, HSP90AB1, UBE2K, EPO* and *UBE2G1.* Quantitative PCR was performed to validate the NGS data consistency - the up-regulated expression of *UBE2B, ATF2, AKT1, NFKBIA, UBE2J1, HSP90B1* along with decreased expression of *NKFBIB, SUMO1, EP300, UBE2G1, UBE2K, HSP90AA1* were confirmed with a comparable level of fold changes (Fig. 3F). Moreover, differentially expressed genes were used to perform protein-protein interaction analysis for the hypoxia signaling pathway with standard parameter settings and predominant interactions were observed in most of the proteins and less interaction was seen for fewer proteins (Figure S2).

### Differentially expressed genes in nitric oxide signaling and mitochondrial function during iPSC-derived cardiomyocyte differentiation

Nitric oxide (NO) is an important molecule involved in several physiological and pathological mechanisms in humans (43). Under normal conditions, NO is produced and acts as a messenger and cytoprotective antioxidant (43, 44). During iPSC induced cardiomyocyte differentiation, about 180 genes are detected in nitric oxide signaling pathway, in which 78 genes are significantly increased and 48 genes are down-regulated (Figure 4A-B). From this list, the top 15 up and down-regulated genes are used to generate the heat-maps. An extensive list of genes is increased during cardiomyocyte differentiation including, *GUCY1A1, RHOJ, LYZ, TLR*, and *PPP1R14C; PPM1J, MAP3K15, PRKCB, PIK3R6* and *PIK3R5* are decreased during cardiomyocyte differentiation. Quantitative PCR experiments were performed to validate the consistency of RNA sequencing and *RHOU, APOD, TLR4 PDGFC, VEGFD, CACNA1C* gene expression were augmented, while *NOS2, FLT4, PIK3CD, PPM1J PED2A, MAP3K15* were down-regulated in differentiated cardiomyocytes confirming the sequencing data (Figure 4C). Furthermore, to investigate the interactivity of significantly altered genes in nitric oxide signaling, we performed protein-protein interaction analysis with standard parameter settings. Strong and multiple interactions were observed between most of the proteins and less or no interactions were seen with fewer proteins (Figure S3).

**Figure 4.**
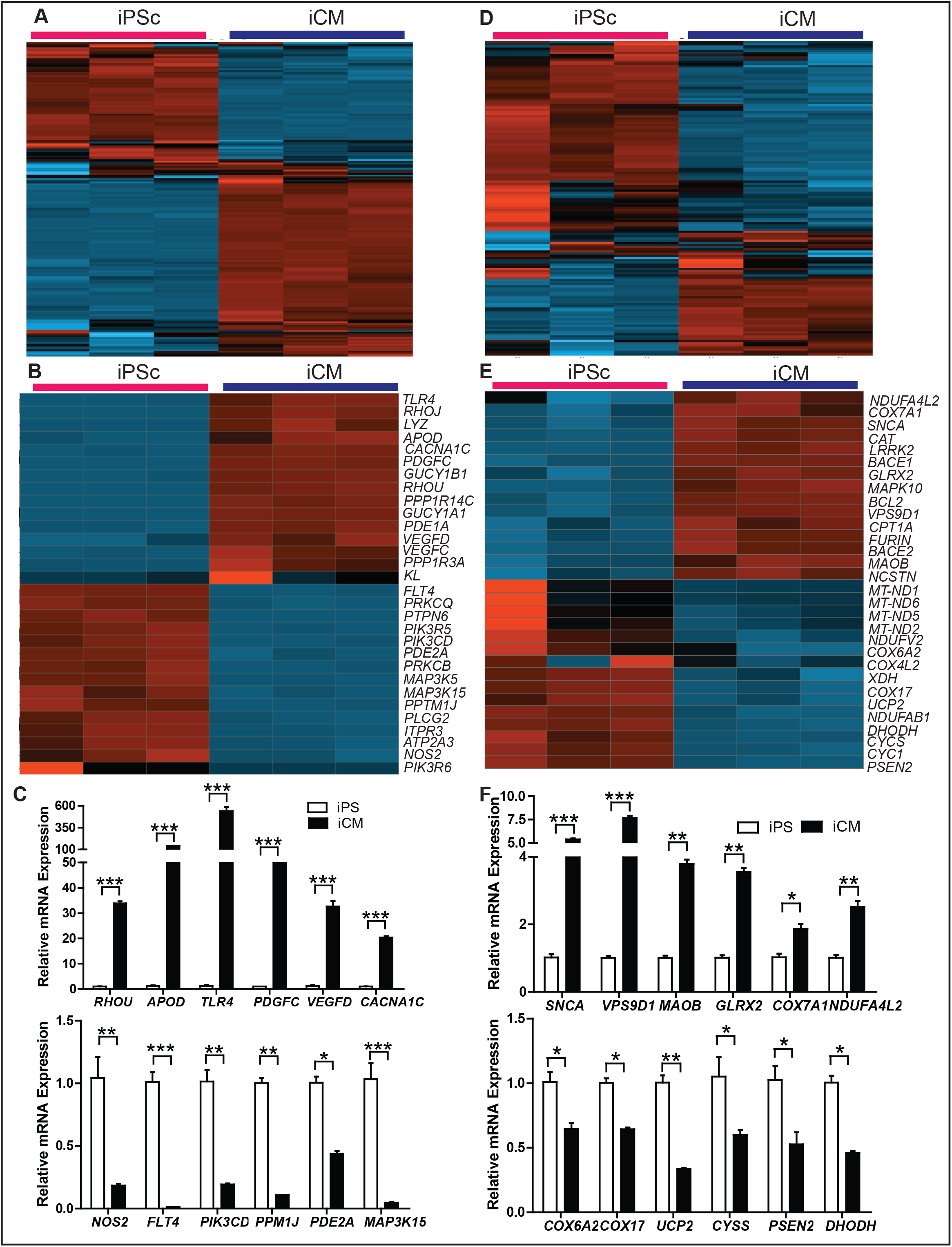
Differentially altered Canonical pathways in biological relevance with Redox signaling: A and D) Heat map representation of all genes involved in nitric oxide signaling and mitochondrial functions respectively identified using IPA analysis from RNA-seq of *i*PSs vs *i*CMs cells. **B and E)**. Heat map illustrating top 15 up and down regulated DEGs in nitric oxide signaling and mitochondrial functions respectively identified by IPA using RNA-Seq. **C and F)**. qPCR validation of 6 up and down regulated genes from each respective pathway validating the significant changes in the expression when compared between *i*PSs vs *i*CMs cells. Gapdh was used as control (n=4-6/group). *p<0.05, **p<0.001, ***p<0.0001.

Mitochondria are the main source of energy production through the process of respiration and majorly involved in production of reactive oxygen species (45). Recent studies revealed that mitochondrial fusion was crucial for differentiation of iPSC cells into cardiomyocytes (46). During iPSC induced cardiomyocyte differentiation, 144 genes involved in mitochondrial functions were detected. Among them, 21 genes were significantly increased and 37 genes were down-regulated (Figure 4D-E). From this list, up and down-regulated genes in each (15) were used to generate the heat-maps. The following list of genes were highly increased during induced cardiomyocyte differentiation including, *LRRK2, MAPK10, BCL2, BACE and COX7A1; MT-ND1, DHODH, UCP2, MT-ND6* and *COX6A2* were down-regulated during cardiomyocyte differentiation. To check the RNA-seq data for variability, qPCR analysis were performed and *SNCA, VPS9D1, MAOB, GLRX2, COX7A1, NDUFA4L2* gene expression were highly expressed, however *COX6A2, COX17, UCP2, CYSS, PSEN2, DHODH* were down-regulated supporting the sequencing data in *i*PSC derived cardiomyocytes (Figure 4F). Additionally, to understand the genes involved in regulating mitochondrial functions and their encoded proteins during cardiomyocyte development, we performed STRING analysis with standard parameter settings. The outcome of the STRING analysis showed several strong interactions between half of the proteins; less interaction was seen between fewer proteins, and no interactions were seen between three proteins. (Figure S4).

### Transcriptional expression analysis for antioxidant action of vitamin C and protein ubiquitination transcriptomes during iPS-cardiomyocyte differentiation

Several exogenous and endogenous antioxidants play inhibitory or compensatory roles against ROS signaling (47) including Vitamin-C (Ascorbate or Ascorbic Acid), Tocopherols, Glutathione, and Superoxide Dismutase, Catalases and Peroxidases (48). Most of these antioxidants are endogenously regulated by NFE2L2/Nrf2, as described earlier. Here we are demonstrating the role of the genes that are involved in vitamin C antioxidant function (49). Endogenous activation and exogenous supplementation of Vitamin C scavenging ROS and reactive nitrogen species combines with Vitamin E and protects both hydrophobic compounds including membrane lipids (49). During cardiomyocyte differentiation, about 65 genes were detected in antioxidant action of Vitamin C pathway, in which 21 genes were significantly up-regulated and 19 genes were down-regulated (Figure 5A-B). From this list, the top 15 up and down-regulated genes were used to generate the heat-maps. An extensive list of genes were increased during cardiomyocyte differentiation including, *PLCL1, PLA2G4A, PLD1, PLD4, PLA2R1.* Further *SLC2A3, PLCG2, PLBD1, PLA2G16 and PLA2G7* were decreased during cardiomyocyte differentiation. Quantitative PCR experiments were performed to check the variability of RNA sequencing showing that *PLD4, JAK2, RELA, PLCL1, NFKB1, PNPLA8* gene expression were upregulated in cardiomyocyte differentiation while *GPLD1, STAT5A,PLD2, HMOX1, PLBD1, PLCL2* were down-regulated in differentiated cardiomyocytes compared to undifferentiated iPSC cells confirming the sequencing data (Figure 5C). To investigate the interactions of genes involved in regulating antioxidant action of vitamin C during cardiomyocyte differentiation, we performed STRING analysis with standard parameter settings. The STRING analysis showed several strong interactions between most of the proteins and less interaction between fewer proteins. In this case, more than ten proteins didn’t show any interactions. (Figure S5).

**Figure 5.**
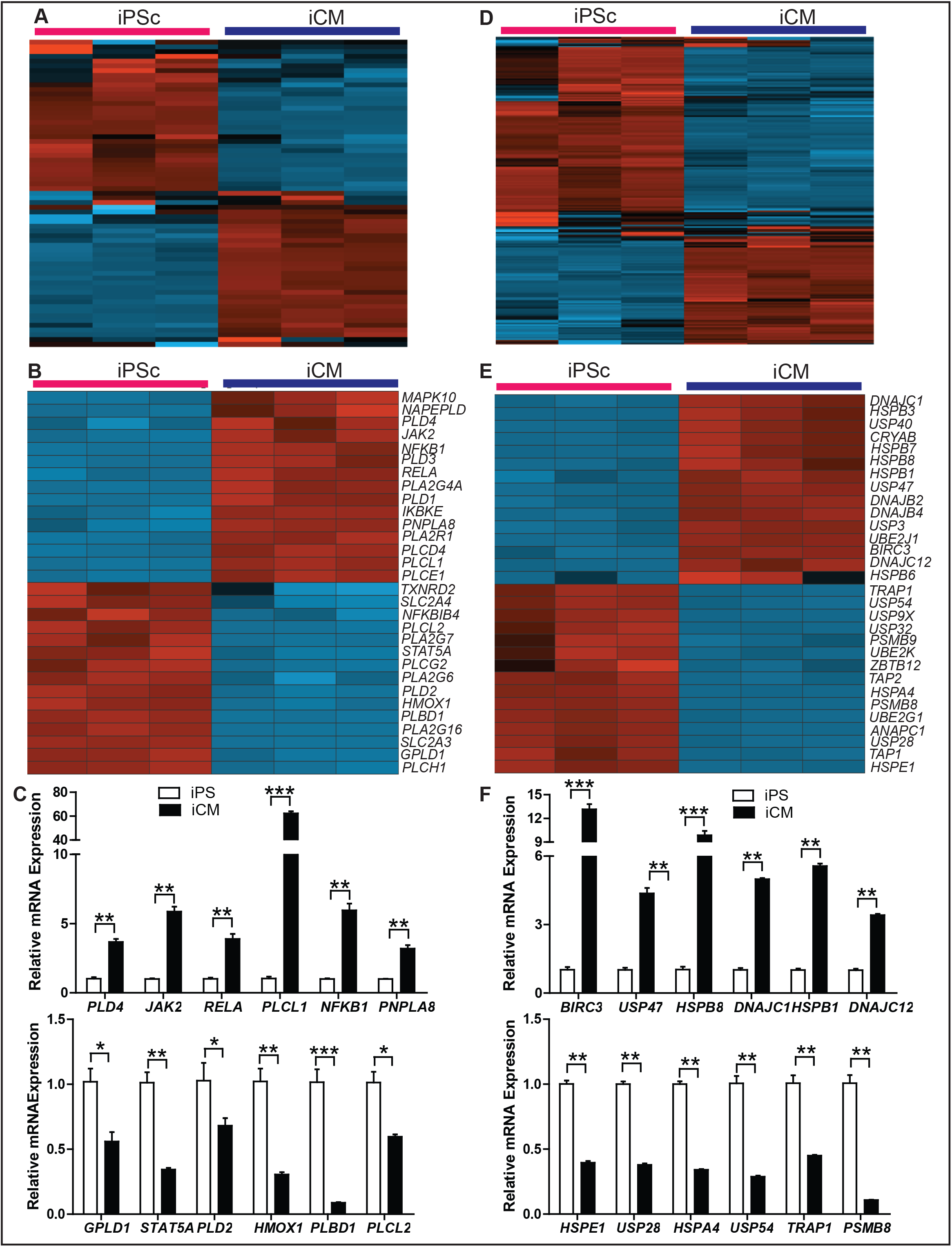
Evaluation of Vitamin C Antioxidant signaling and Protein Ubiquitination Statuses of iCMs vs iPSCs: A and D) Heat map of the genes involved in antioxidant action of Vitamin C and protein ubiquitination mechanisms identified from RNA-seq of *i*PSs and *i*CMs cells. B and E). Top 15 up and down regulated DEGs in antioxidant action of Vitamin C and protein ubiquitination mechanisms represented by heat maps. **C and F)**. qPCR validation of 6 up and down regulated genes from each respective pathway validated the significant changes in the expression compared between *i*PSs vs *i*CMs cells. Gapdh was used as control (n=4-6/group). *p<0.05, **p<0.001, ***p<0.0001.

The ubiquitin proteasome pathway is required for targeted degradation of most short-lived proteins in humans (50). Ubiquitination is a process in protein metabolism and ubiquitination of proteins involves attaching ubiquitin to a specific protein followed by its inactivation and targeting it to a to degradation pathway (51). During iPS-derived cardiomyocyte differentiation, a large number of transcriptional factors involved in reprogramming the iPSC cells into cardiomyocytes is seen, in which lots of proteins (stable and short lived) are synthesized and require degrading or metabolizing in a timely manner to avoid unfolded response or accumulation of misfolded proteins (52). In RNA-sequencing of iCMs about 217 genes were detected, which are involved in protein ubiquitination pathway, 40 genes were highly expressed and 65 genes were significantly down-regulated (Figure 5D-E). From this list, the up and down-regulated genes in each (15 genes) were used to generate the heat-maps.

The following list of genes were increased highly during induced cardiomyocyte differentiation including, *HSPB3, HSPB7, CRYAB, BIRC3 and HSPB6; TAP1, TAP2, USP9X, PSMB9, and PSMB8* were greatly down-regulated during cardiomyocyte differentiation. Furthermore, to verify and validate the RNA-seq data for consistency, qPCR analysis was performed showing that *BIRC3, USP47, HSPB8, DNAJC1,HSPB1* and *DNAJC12* genes were highly expressed, and *HSPE1, USP28, HSPA4, USP54, TRAP1 and PSMB8* were down-regulated recognizing the consistency of the sequencing data in *i*PSC derived cardiomyocytes (Figure 5F). To investigate the interactions of genes and proteins involved in regulating protein ubiquitination during cardiomyocyte differentiation, STRING analyses were performed with standard parameter settings. The results of analysis showed very strong interactions between majorities of proteins involved in this pathway, whereas seven proteins didn’t involve in any interactions. (Figure S6).

## Discussion

Unlimited potential of iPSC cells for proliferation and differentiation into various lineages has a clinical significance to treat diseases associated with cellular and organ damage (8, 22). In the past decade, differentiation of iPSCs to ventricular cardiomyocytes has been an attractive source of cardiac regenerative therapy (1, 53, 54). These induced cardiomyocytes also provide a reliable platform for researchers to study the disease mechanisms, drug screening and gene specific gain and loss of functions (6-9) at the cellular level. Recently, these differentiated cardiomyocytes have been largely used to treat injured myocardium (I/R or MI injury) to support the regeneration and regain the myocardial structure and function (55-57). However, the rate of repair is not remarkable possibly due to reduced survival and poor engraftment. Previous transcriptome studies focused only on the genes involved in differentiation/reprogramming, but failed to show the essential mechanisms required for survival of these cells against stress at the implantation site (injury site).

In the current study, we used next generation RNA sequencing and performed transcriptome profiling during differentiation of iPSCs into iCM cells. The global gene expression profiling, qPCR validation, IPA and gene ontology analysis confirmed that expression patterns became more specified towards cardiac specific differentiation and cardiac developmental networks as described widely (1, 2, 21, 36). Pathway analysis for highly expressed genes (>50- and > 100-fold changes) showed a clear description of changes related to cardiac reprogramming and development related changes (Figure 1 and 2). Further, we found that the 5 to 20-fold change in transcripts involved key redox signaling pathways including Nrf2 mediated glutathione signaling, hypoxia and nitric oxide signaling, mitochondrial functions and ubiquitination most of them were differentially expressed. Considering the sensitivity and susceptibility of these differentiated cardiomyocytes to exogenous stressors (oxidative, inflammatory, fibrotic), and the cross talk between cardiomyocytes survival and potential of redox signaling mechanisms could be novel.

Nrf2 mediated glutathione signaling plays an essential role in protection against pathological maladaptation by suppressing ROS levels and oxidant stress (33, 37, 39). Glutathione is a small, key thiol molecule that is used to scavenge ROS levels in the living system and protect against the cellular toxicity (39). Interestingly, differentiated cardiomyocytes displayed a decreased expression of *GSTM5, AOX1* levels which leads to possible increase in reactive oxygen species levels as these genes are involved in the regulation of ROS homeostasis (58). Further, *MGST1* is confined in the endoplasmic reticulum (ER) and outer mitochondrial membrane to protect from oxidant stress, the decreased expression of this gene leads to susceptible for ROS mediated stress in the ER resulting in ER stress (59). However, the increased expression of *GSTA3, GSTM3, GSTM2, CAT* and *NQO1* could be a compensatory response to ROS in these cells during differentiation of cardiomyocytes and provide cyto-protection. Though these transcripts levels are increased, the glutathione metabolizing enzymes *GCLC, GCLM, GPX family enzymes (GPX1, GPX3, GPX4)* and *GSR* are significantly decreased which leads to reduced levels of glutathione availability for scavenging the ROS stress in these differentiated cardiomyocytes. (Figure 3).

Low levels of oxygen induce hypoxic condition; several hypoxia-inducible factors can be activated to protect the system from hypoxic stress. Hypoxic signaling naturally occurs during developmental stages and helps cellular differentiation (29, 40, 42). However, chronic hypoxia condition leads to chemo-resistance in cancer and pathogenesis in several organs (42, 60). During hiCMs differentiation, we observed a decreased expression of *EP300, HSP90AA1* and *HSP90AB1* genes resulting in reduced hypoxic responses and decreased wound healing activity (61). As stable or increased NOS3 protects from chronic hypoxia in embryonic developmental stages (62), decreased *NOS3* expression was seen in differentiated cardiomyocytes which leads to apoptosis in these cells (62). Further, chronic hypoxia induces insulin resistance in cells (63); Tan et al showed that activation of *SLC2A4* induces GLUT4 and protects from insulin resistance in rats (64), decreased *SLC2A4* expression during *i*CM differentiation possibly reduces GLUT4 and induces insulin resistance in these cells (Figure 3).

Nitric oxide (NO) is a fundamental signaling molecule involved in several physiological processes including host defense, and the regulation of vascular tone as well as a potent cytotoxic effector involved in human diseases pathogenesis (62), Griffin et al showed that decreased *ATP2A3* expression induces unfolded protein response (UPR) in Jurkat cells (65) and our sequencing results also observed that down-regulation of *ATP2A3* resulted in induction of unfolded protein response in these cells. Recently, researchers showed decreased *PDE2A* expression leads to increased accumulation of cAMP and myocardial contractility after beta-adrenogenic receptor stimulation (66, 67). Sequencing results also showed down-regulation of *PDE2A* level. Further increased *MAP3K15* activates apoptosis signaling, however iCM cells showed decreased *MAP3K15* levels which suggest the decreased possibility for the apoptosis in these cells under unstressed conditions (Figure 4).

Mitochondria are the powerhouse of cells which synthesize proteins for electron transport and oxidative phosphorylation, to help cellular respiration (68). In oxidative phosphorylation, ATP is generated within the mitochondria and used as an energy source. *UCP2* is one of the major proteins in the mitochondrial anion carrier protein family, which facilitates transfer of anions from the inner to the outer mitochondrial membrane and reduces mitochondrial membrane potential (69). Decreased expression of *UCP2* during iCM cell differentiation leads to possible increase in mitochondrial membrane potential, which could impact the cellular survival in these differentiated cells as previously, Zorova et al showed increased mitochondrial membrane potential could decrease cell survival (70). Further, *COX6A* is the major complex IV enzyme in oxidative phosphorylation and decreased or knockdown of this gene in mouse model showed elevated formation of reactive oxygen species (71). Likewise, in our study the *COX6A* gene expression was decreased during cardiomyocyte differentiation, which might induce apoptosis and necrosis in cells (71). *COX7A1* is expressed predominantly in heart and skeletal muscle (72); Previously Hwang et al showed that increased *COX7A1* expression was observed in hypoxic condition (73), similarly during *i*CMs differentiation also *COX7A1* expression is *augmented* in which might be a compensatory response for hypoxic condition (Figure 4).

Vitamin C or ascorbic acid regulates cell proliferation, differentiation and plays a major role in detoxifying the body from heavy metals (47). Vitamin C is also involving in collagen synthesis, wound healing activities and act as a free radical scavenging antioxidants (13, 48). Guaiquil et al reported that vitamin C protected the myocardium from ischemic injury and decreases the ROS levels and inhibited the activation of Ser/Thr and Tyr protein phosphatases including Phospholipase-C (PLC), PLD (Phospholipase-D), and PLA2 (Phospholipase-A2) to loss/disassembly of tight and adherent junctions in the injured (MI) myocardium (74). Our results in differentiated cardiomyocytes also showed increased *PLCL1, PLA2G4A, PLD1, PLD4, PLA2R1*, and *PLCD4.* Further, most of the phospholipase genes were increased during differentiation and few of them including *PLCG2, PLBD1, PLA2G16, PLA2G7, HMOX1* and TXNRD2 were decreased. These results suggest that the down regulation of these genes ultimately induces ROS generation and oxidative stress (Figure 5).

Overall, the current study identified that new candidate genes and pathways that are unidentified in previous studies significantly changed during cardiomyocyte differentiation compared with induced iPSC cells. Specifically, we have showed that the pathways and genes that are involved in redox signaling are highly differentially regulated and significantly responsible for ROS scavenging effects. The successful maintenance of redox signaling in differentiated cardiomyocytes will provide a new insight for devising mechanism-based in vivo cell survival strategies during cell therapy.

**Table-1:** Top altered canonical Pathways and its significance up to 10 fold Changes.

| **IPA No** | **Top altered canonical Pathways** | **>100** | **50-100** | **10-50** |
| --- | --- | --- | --- | --- |
| 1 | Colorectal Cancer Metastasis Signaling | -3.61 | -2.00 | -3.13 |
| 2 | ILK Signaling | -3.16 | -2.00 | -1.81 |
| 3 | Glioblastoma Multiforme Signaling | -3.00 | -1.34 | -4.00 |
| 4 | GP6 Signaling Pathway | -2.89 | -1.13 | -1.60 |
| 5 | Signaling by Rho Family GTPases | -2.89 | -1.00 | -1.96 |
| 6 | Phospholipase C Signaling | -2.83 | -1.00 | -3.41 |
| 7 | IL-8 Signaling | -2.83 | -1.00 | -3.41 |
| 8 | PCP pathway | -2.83 | -0.82 | -2.31 |
| 9 | HMGB1 Signaling | -2.83 | -0.82 | -1.27 |
| 10 | Cardiac Hypertrophy Signaling | -2.68 | -0.45 | 0.00 |
| 21 | Production of Nitric Oxide and Reactive Oxygen Species in Macrophages | -2.24 | 1.00 | 0.40 |
|  |  | **10-50** | **5-10** | **2-5** |
| 169 | Antioxidant Action of Vitamin C | 0.45 | 0.91 | 0.54 |
| 188 | NRF2-mediated Oxidative Stress Response | -1.13 | 0.00 | 0.21 |
| 201 | Glutathione Redox Reactions I | - | -1.00 | - |

## Acknowledgments

This study was supported by funding from NHLBI (2HL118067), NIA (AG042860), the AHA (BGIA 0865015F), University of Utah Center for Aging Pilot grant (2009), the Division of Cardiovascular Medicine/Department of Medicine, University of Utah and the start-up funds (for NSR) by Department of Pathology, the University of Alabama at Birmingham, AL and UABAMC21 grant by the University of Alabama at Birmingham, AL.

Authors thank Dr. Michael Crowley, Director, Heflin Center for Genomic Sciences for providing access to the genomic core facilities. Authors also thank Dr. Anilkumar Challa for editorial assistance.

