## Supplemental Table and Figures for "Impaired Regulation of Redox Transcriptome during the Differentiation of *i*PSCs into *Induced* Cardiomyocytes (*i*CMs)"

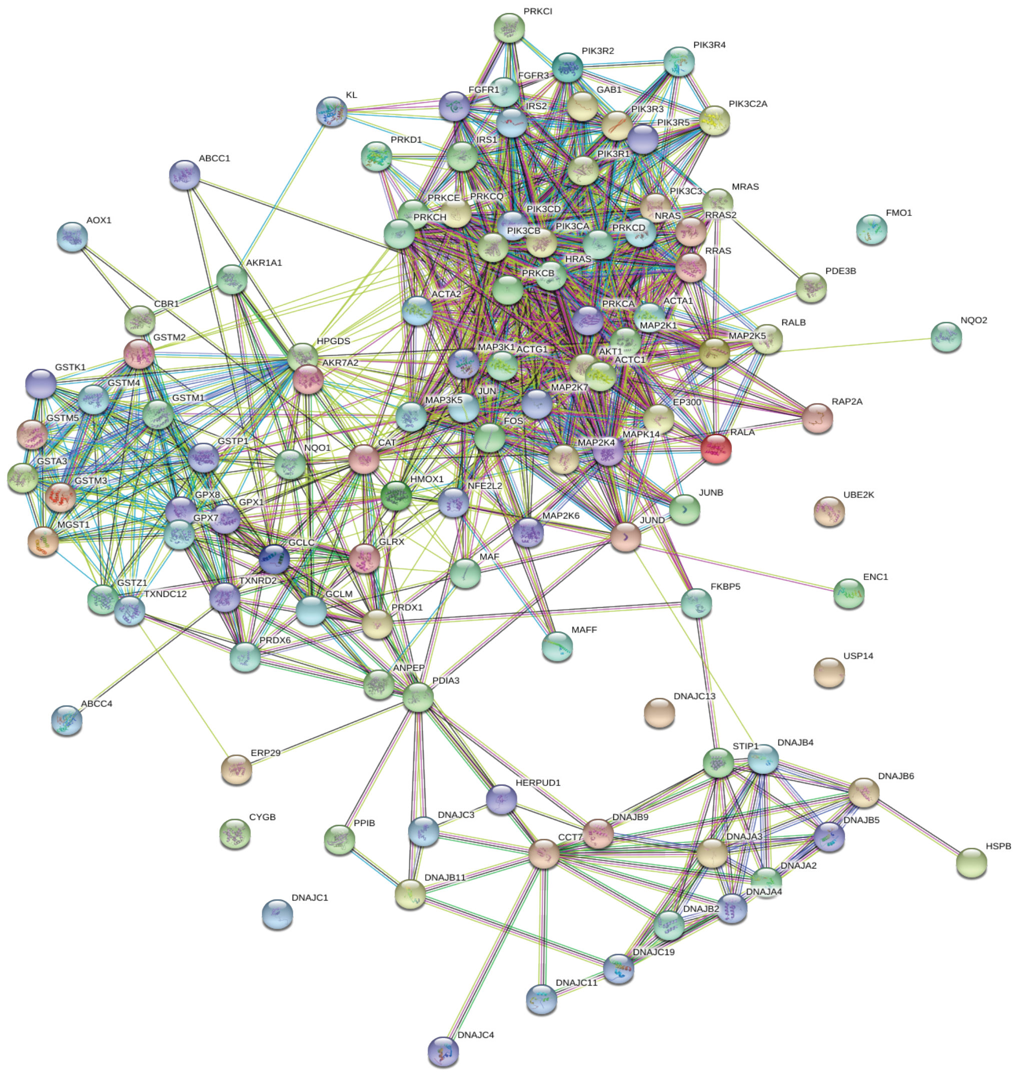


**Figure S1. Protein interaction networks for Nrf2 mediated oxidative response pathway.** Protein-protein interaction network based on genes (114 genes) that are involved in Nrf2 mediated oxidative response pathway which are significantly (1.5-fold and above with p>0.05) changed in hiCMs when compared with undifferentiated hiPSC cells. A higher interactivity was seen among proteins that are changed in hiCMs. Color nodes represents the first shell of interactors and white node describes second shell of interactors. Node with shapes defines the availability of the 3D structures and connecting blue line indicates predicted interactions and pink/purple connecting lines indicates the experimental proved interactions. Yellow line represents the text-mining between protein interactions.

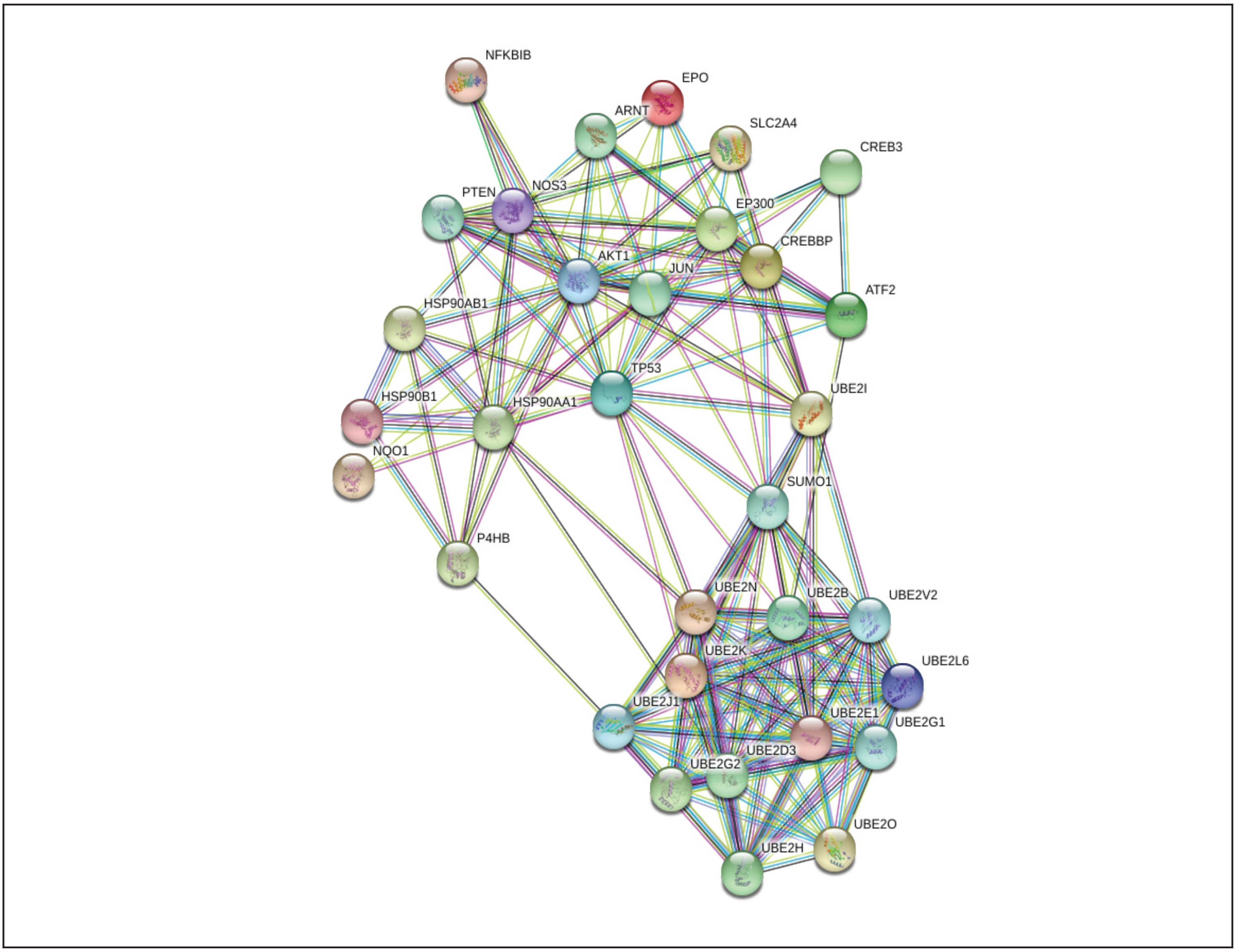


**Figure S2. Protein interaction networks for hypoxia signaling transcripts.** Protein-protein interaction network based on genes (29 transcripts) that are involved in hypoxia signaling which were significantly (1.5-fold and above with p>0.05) changed in hiCMs when compared with undifferentiated hiPSC cells. A higher interactivity was seen among proteins that are changed in hiCMs. Node with shapes expresses the 3D structures and connecting blue line indicates predicted interactions and pink/ purple connecting lines indicates the experimental evidenced interactions. Color nodes represent the first shell of interactors and white node describes second shell of interactors. Yellow line indicates the text-mining between protein interactions.

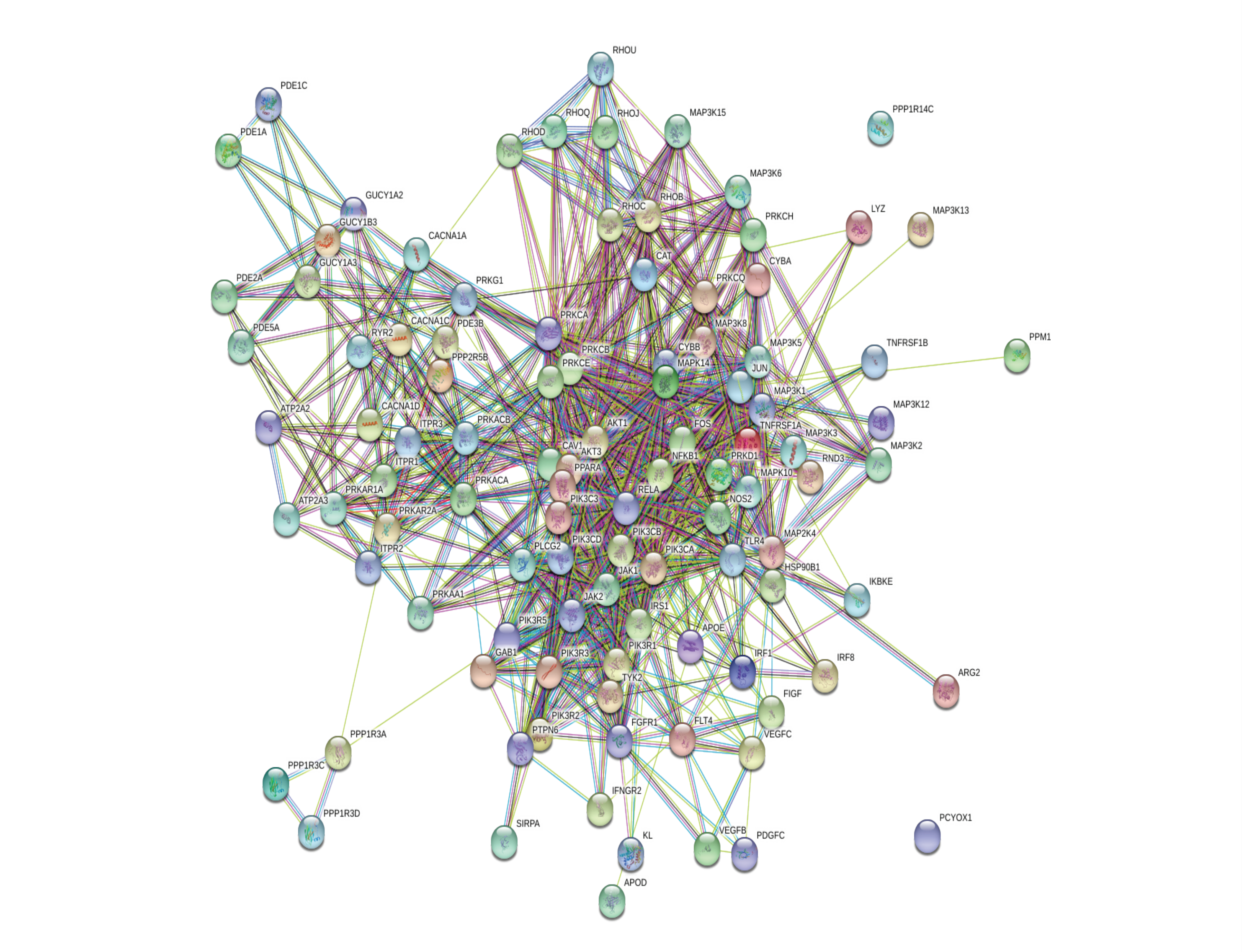


**Figure S3. Protein interaction networks for Nitric Oxide Signaling in the Cardiovascular System.** Protein-protein interaction networks for genes (126 genes) significantly (1.5-fold and above with p>0.05) changed that are involved in Nitric Oxide Signaling (NOS) pathway in hiCMs when compared with undifferentiated hiPSC cells. A higher interactivity is seen among proteins that are changed in hiCMs. Color nodes signify the first shell of interactors and white node describes second shell of interactors. Node with shapes expresses the available 3D structures and blue connecting line indicates predicted interactions and pink/purple connecting lines indicates the experimental demonstrated interactions. Yellow line represents the text-mining between protein interactions.

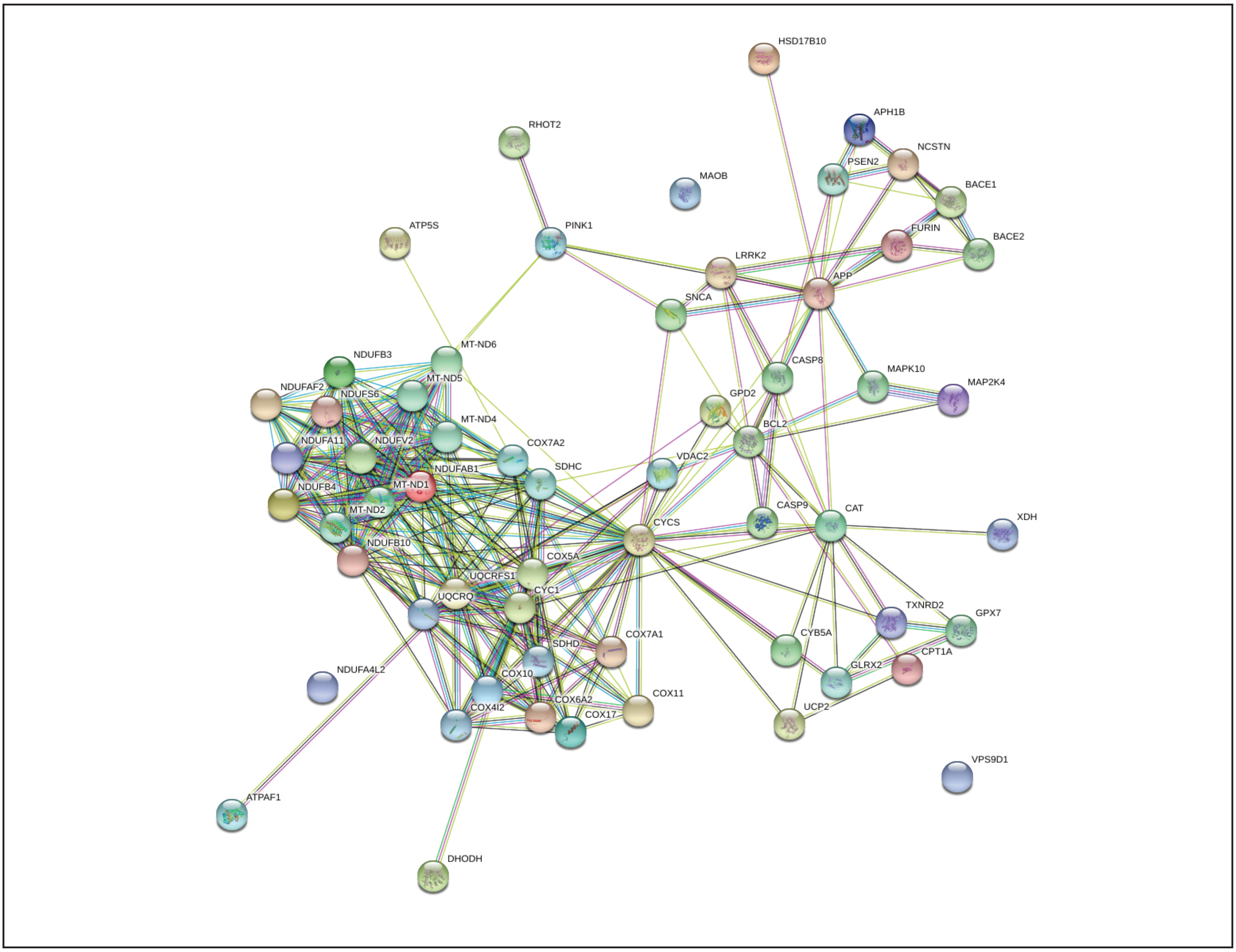


**Figure S4. Protein interaction networks for mitochondrial functions.** Protein-protein interaction network based on genes (58 transcripts) that are involved in mitochondrial functions which are significantly (1.5-fold and above with p>0.05) changed in hiCMs differentiation when compared with undifferentiated hiPSC cells. A higher interactivity was seen among proteins that are changed in hiCMs. Node with shapes expresses the 3D structures and connecting blue line indicates predicted interactions and pink/purple connecting lines indicates the experimental evidenced interactions. Color nodes represent the first shell of interactors and white node describes second shell of interactors. Yellow line indicates the text-mining between protein interactions.

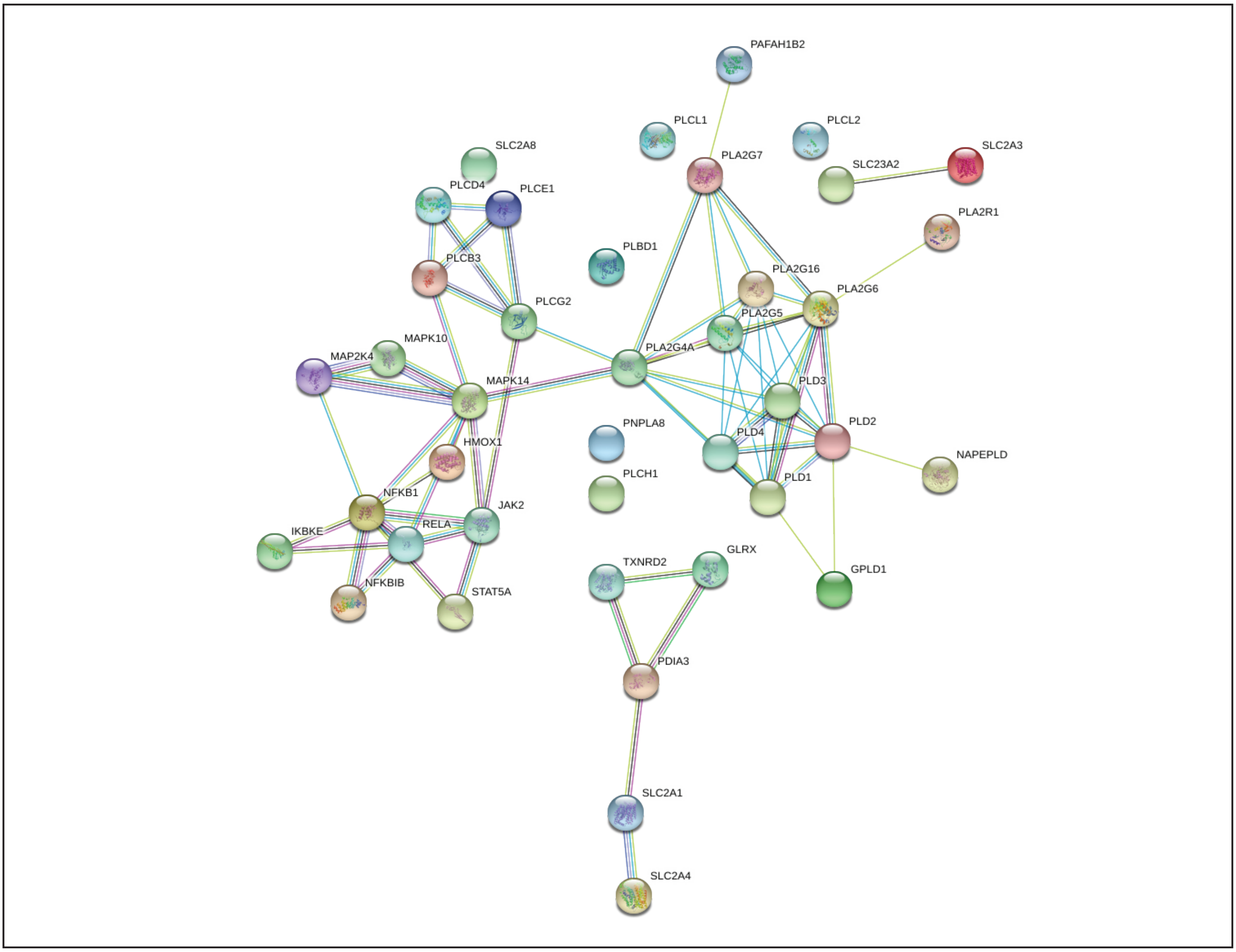


**Figure S5. Figure S5. Protein interaction networks for antioxidant action of Vitamin C pathway.** Protein-protein interaction network based on genes (40 genes) that are involved in antioxidant action of Vitamin C which were significantly (1.5-fold and above with p>0.05) changed in hiCMs when compared with undifferentiated hiPSC cells. A higher interactivity was seen among proteins that are changed in hiCMs. Color nodes represents the first shell of interactors and white node describes second shell of interactors. Node with shapes defines the availability of the 3D structures and connecting blue line indicates predicted interactions and pink/purple connecting lines indicates the experimental proved interactions. Yellow line represents the text-mining between protein interactions.

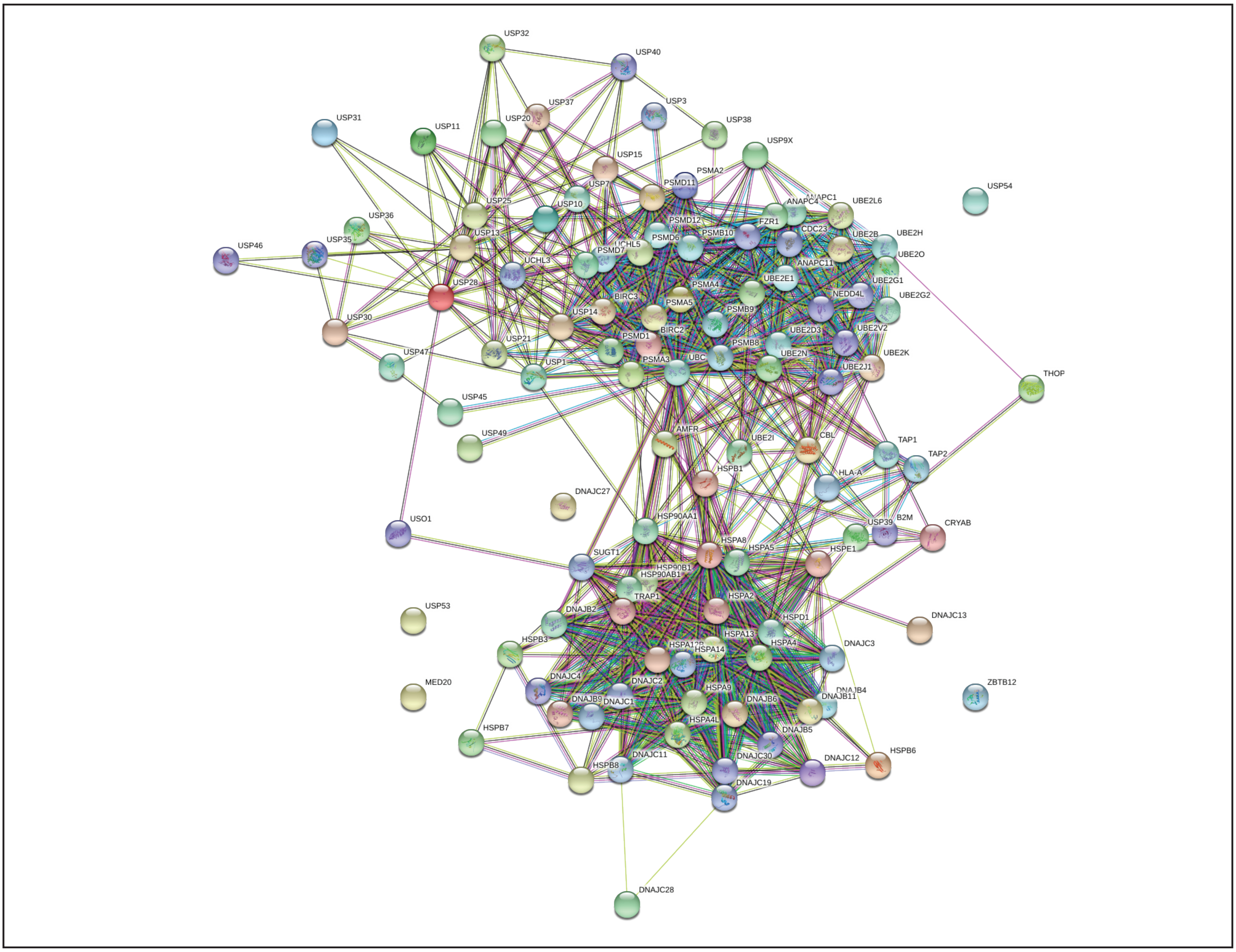


**Figure S6. Protein interaction networks for protein ubiquitination transcripts.** Protein-protein interaction network based on genes (105 transcripts) that are involved in protein ubiquitination which are significantly (1.5-fold and above with p>0.05) changed in hiCMs when compared with undifferentiated hiPSC cells. A higher interactivity was seen among proteins that are changed in hiCMs. Node with shapes expresses the 3D structures and connecting blue line indicates predicted interactions and pink/purple connecting lines indicates the experimental evidenced interactions. Color nodes represent the first shell of interactors and white node describes second shell of interactors. Yellow line indicates the text-mining between protein interactions.

Supplemental Table-1: List of Real-Time qPCR Primer Sequences.

| ***Gene Name*** | **Forward Primer** | **Reverse Primer** |
|---|---|---|
| *GSTA3* | *CCTTGAGAAGCTGAGCGGAG* | *CATTCTGCCCCGTCCATTGAA* |
| *ACTA2* | *CCGGGACTAAGACGGGAATC* | *TTGTCACACACCAAGGCAGT* |
| *GSTM3* | *TCGAGTGGACATCATAGAGAACC* | *TGGTCAGAGCTGTAACAGAGC* |
| *JUNB* | *ACAAACTCCTGAAACCGAGCC* | *CGAGCCCTGACCAGAAAAGTA* |
| *PRKD1* | *CTTTTTCGCCATGACCCTACC* | *GGAAGCTGACAAGACCACTTCA* |
| *MARS* | *CAAGGACCCACCTCTCAATGT* | *TCTCCCCGCCATTTGGTTTTC* |
| *NQO2* | *CCACGAAGCCTACAAGCAAAG* | *CCAGTACAGCGGGAACTGAAATA* |
| *AOX1* | *ATGCCTGTCTGATTCCCATCT* | *CATGACACTTGGCAATCCTCT* |
| *FGFR1* | *CCCGTAGCTCCATATTGGACA* | *TTTGCCATTTTTCAACCAGCG* |
| *MGST1* | *ATTGGCCTCCTGTATTCCTTGA* | *GTGCTCCGACAAATAGTCTGAAG* |
| *GSTM5* | *CCATCCTGCGCTACATTGC* | *CCAGCTCCATGTGGTTATCCAT* |
| *PRKCQ* | *AGAACGGGCAGATGTATATCCA* | *GGTCCACGTTTTTGCCTTTCA* |
| *UBE2B* | *TCTCTGCTGGATGAACCGAAT* | *ACAATGGCCGAAACTCTTTTCTC* |
| *ATF2* | *AATTGAGGAGCCTTCTGTTGTAG* | *CATCACTGGTAGTAGACTCTGGG* |
| *AKT1* | *AGCGACGTGGCTATTGTGAAG* | *GCCATCATTCTTGAGGAGGAAGT* |
| *NFKBIA* | *CCCTACACCTTGCCTGTGAG* | *TAGACACGTGTGGCCATTGT* |
| *UBE2J1* | *GAGACCCGCTACAACCTGAAG* | *CGCATGGTAATGATCTGTTGGAT* |
| *HSP90B1* | *CCGGTGTAGGAATGACCAGAG* | *TTAAAAACTCGCTTGTCCCAGAT* |
| *NFKBIB* | *CGACACCTACCTCGCTCAG* | *GTCGGAATCGGGGTACAAGG* |
| *SUMO1* | *TGACCAGGAGGCAAAACCTTC* | *AATTCATTGGAACACCCTGTCTT* |
| *EP300* | *AGCCAAGCGGCCTAAACTC* | *TCACCACCATTGGTTAGTCCC* |
| *UBE2G1* | *AGGTGGTGTTTTTAAGGCTCATC* | *CATTTGGGTGCCAGATTTCTGTA* |
| *UBE2K* | *GTTCCGTCACAGGGGCTATTT* | *AATACCGTGCGGAGAGTCATT* |
| *HSP90AA1* | *GCTTGACCAATGACTGGGAAG* | *AGCTCCTCACAGTTATCCATGA* |
| *RHOU* | *TTCCAGAACGTCAGTGAGAAATG* | *GGCTGAACACTCGATGTAGGAG* |
| *APOD* | *ACAAGCATTTCATCTTGGGAAGT* | *CATCAGCTCTCAACTCCTGGT* |
| *TLR4* | *AGACCTGTCCCTGAACCCTAT* | *CGATGGACTTCTAAACCAGCCA* |
| *PDGFC* | *GACTCAGGCGGAATCCAACC* | *CTTGGGCTGTGAATACTTCCATT* |
| *VEGFD* | *ATGGACCAGTGAAGCGATCAT* | *GTTCCTCCAAACTAGAAGCAGC* |

Supplemental Table-1: List of Real-Time qPCR Primer Sequences.

| ***Gene Name*** | **Forward Primer** | **Reverse Primer** |
|---|---|---|
| *CACNA1C* | *TGATTCCAACGCCACCAATTC* | *GAGGAGTCCATAGGCGATTACT* |
| *NOS2* | *TTCAGTATCACAACCTCAGCAAG* | *TGGACCTGCAAGTTAAAATCCC* |
| *FLT4* | *TGCACGAGGTACATGCCAAC* | *GCTGCTCAAAGTCTCTCACGAA* |
| *PIK3CD* | *AAGGAGGAGAATCAGAGCGTT* | *GAAGAGCGGCTCATACTGGG* |
| *PPM1J* | *GGCAAGAGTCGGCACAATGA* | *CTGCTCTCGGATATGGCGATG* |
| *PDE2A* | *GACCGCAAGATCCTCCAACTG* | *CCGAGCACTTTGTCTCCGA* |
| *MAP3K15* | *CCTTCTACGACGCAGATGTTG* | *GCATCGGTGTCATGGTACAAGA* |
| *SNCA* | *AAGAGGGTGTTCTCTATGTAGGC* | *GCTCCTCCAACATTTGTCACTT* |
| *VPS9D1* | *AAGGACAGCTCGTTCGAGGA* | *AGCAGCCTGTCTACGGCATT* |
| *MAOB* | *GGAGCTAGGATTGGAGACCTAC* | *CCCTGAAGGGGTATGATTTGC* |
| *GLRX2* | *TCTTTGGAGAATTTAGCGACGG* | *CTGGTTTCCATATTCAAGCAGGT* |
| *COX7A1* | *CACGGCCTTATGCGTTTCCT* | *TTTTGGAGATTTCCGCCGGG* |
| *NDUFA4L2* | *AGGCTTTGGGCCTGGAATTG* | *CCTGAGTGGGGAGGGATCAG* |
| *COX6A2* | *CCTTCAACTCCTATCTCCACTCG* | *GTTGGTAGGGACGGAACTCG* |
| *COX17* | *TGCGTGTATCATCGAGAAAGGA* | *GCCTCAATTAGATGTCCACAGTG* |
| *UCP2* | *GTCATCGCCTCCCCTGTAGA* | *GAGCATGGTAAGGGCACAGTG* |
| *CYSS* | *TGAGTAATAATTGGCCACTGCCTT* | *AGTTTTAAATCAGGACTGCCCAACA* |
| *PESN2* | *CTGACCGCTATGTCTGTAGTGG* | *CTTCGCTCCGTATTTGAGGGT* |
| *DHODH* | *GTTCTGGGCCATAAATTCCGA* | *TCTGGGTCTAGGGTTTCCTTC* |
| *PLD4* | *GCTTGTCCTTGTGGAAAGCAT* | *AGGGACCAGTAGTATGAAGCC* |
| *JAK2* | *ATCCACCCAACCATGTCTTCC* | *ATTCCATGCCGATAGGCTCTG* |
| *RELA* | *GTGGGGACTACGACCTGAATG* | *GGGGCACGATTGTCAAAGATG* |
| *PLCL1* | *AAAGTCCGGCCAAATTCTCG* | *TTTCCGTGTTTTTCCCCAGTC* |
| *NFKB1* | *AACAGAGAGGATTTCGTTTCCG* | *TTTGACCTGAGGGTAAGACTTCT* |
| *PNPLA8* | *CAGCGAGAAAAGATTATCGCAAG* | *AGCTTTGGGTCAGTTGTTCTTC* |
| *GPLD1* | *ATGTCTGCTTTCAGGTTGTGG* | *ACGCATCCTGGTGTTCTAGTAA* |
| *STAT5A* | *CGACGGGACCTTCTTGTTG* | *GTTCCGGGGAGTCAAACTTCC* |
| *PLD2* | *TCGATTTGCCGTTGCCTATTC* | *GGTCAAGAGACGGTTGAGGTA* |
| *HMOX1* | *AAGACTGCGTTCCTGCTCAAC* | *AAAGCCCTACAGCAACTGTCG* |

Supplemental Table-1: List of Real-Time qPCR Primer Sequences.

| ***Gene Name*** | **Forward Primer** | **Reverse Primer** |
|---|---|---|
| *PLBD1* | *GCAACTGCATACTGGATGCCT* | *TCAGGGTTTGAGAGCCATAGC* |
| *PLCL2* | *TTCAGAACTCAAAAAGGTTCGCT* | *GCTGCGGAATATGTCTGTGTT* |
| *BIRC3* | *TTTCCGTGGCTCTTATTCAAACT* | *GCACAGTGGTAGGAACTTCTCAT* |
| *USP47* | *CTCGACGCTAATTTTGAGCCA* | *CTCTTGGAAGCGGACCTATAAAC* |
| *HSPB8* | *GCACAGCTTCAAGCCAGAG* | *CAAATGTTGAGTAAGGAGGGACC* |
| *DNAJC1* | *AAAACAGGAACCGAACACAGAA* | *GGGCAATCTTTTCCCATCGAC* |
| *HSPB1* | *ACGGTCAAGACCAAGGATGG* | *AGCGTGTATTTCCGCGTGA* |
| *HSPE1* | *ATGGCAGGACAAGCGTTTAGA* | *CCGATCCAACAGCGACTACT* |
| *USP28* | *CCCTGGATCTATTAAAGGGAGCA* | *GCTGGAATGCGTCCTCTAGC* |
| *HSPA4* | *GCATCGAGACTATCGCTAATGAG* | *TGCAAGGTTAGATTTTTCTGCCT* |
| *USP54* | *CTGTCCAAGCAACTGTGGAGA* | *CCAATCGTGATAATCTGTGGAGC* |
| *TRAP1* | *CCAATGCCGAGAAAGGCAC* | *CAAACTGGCCGATGATCTTGC* |
| *PSMB8* | *GCAGGCTGTACTATCTGCGAA* | *AGAGCCGAGTCCCATGTTCAT* |
